# GTSF-1 is required for the formation of a functional RNA-dependent RNA Polymerase complex in *C. elegans*

**DOI:** 10.1101/271270

**Authors:** Miguel Vasconcelos Almeida, Sabrina Dietz, Stefan Redl, Emil Karaulanov, Andrea Hildebrandt, Christian Renz, Helle D. Ulrich, Julian König, Falk Butter, René F. Ketting

**Affiliations:** Biology of Non-coding RNA Group, Institute of Molecular Biology, Ackermannweg 4, 55128 Mainz, Germany; Quantitative Proteomics Group, Institute of Molecular Biology, Ackermannweg 4, 55128 Mainz, Germany; Bioinformatics Core Facility, Institute of Molecular Biology, Ackermannweg 4, 55128 Mainz, Germany; Genomic Views of Splicing Regulation Group, Institute of Molecular Biology, Ackermannweg 4, 55128 Mainz, Germany; Maintenance of Genome Stability Group, Institute of Molecular Biology, Ackermannweg 4, 55128 Mainz, Germany

## Abstract

In every domain of life, Argonaute proteins and their associated small RNAs regulate gene expression. Despite great conservation of Argonaute proteins throughout evolution, many proteins acting in small RNA pathways are not widely conserved. Gametocyte-specific factor 1 (Gtsf1) proteins, characterized by two tandem CHHC zinc fingers and an unstructured, acidic C-terminal tail, are conserved in animals and act in small RNA pathways. In fly and mouse, they are required for fertility and have been shown to interact with Piwi clade Argonautes. We identified *T06A10.3* as the *Caenorhabditis elegans* Gtsf1 homolog and named it *gtsf-1*. Given its conserved nature and roles in Piwi-mediated gene silencing, we sought out to characterize GTSF-1 in the context of the small RNA pathways of *C. elegans*. Like its homologs, GTSF-1 is required for normal fertility. Surprisingly, we report that GTSF-1 is not required for Piwi-mediated gene silencing. Instead, *gtsf-1* mutants show strong depletion of a class of endogenous small RNAs, known as 26G-RNAs, and fully phenocopy mutants lacking RRF-3, the RNA-dependent RNA Polymerase that synthesizes 26G-RNAs. We show, both *in vivo* and *in vitro*, that GTSF-1 specifically and robustly interacts with RRF-3 via its tandem CHHC zinc fingers. Furthermore, we demonstrate that GTSF-1 is required for the assembly of a larger RRF-3 and DCR-1-containing complex, also known as ERIC, thereby allowing for 26G-RNA generation. We propose that GTSF-1 homologs may similarly act to drive the assembly of larger complexes that subsequently act in small RNA production and/or in imposing small RNA-mediated silencing activities.

## Introduction

Endogenous small non-coding RNAs are responsible for regulating gene expression in many organisms. These small RNAs (sRNAs) act within the context of RNA interference (RNAi) or RNAi-like pathways. In a variety of situations, these pathways provide an RNA-based protection against “foreign” genetic elements such as transposable elements (TEs) and viruses (Ketting, 2011; Luteijn and Ketting, 2013).

In many RNAi-like pathways, sRNAs are generated from double-stranded RNA (dsRNA) precursors by Dicer, a conserved RNase III-related enzyme (Ketting, 2011). Subsequently, sRNAs associate with Argonaute family proteins, and guide them to target transcripts with complete or partial sequence complementarity. Upon Argonaute binding, transcripts are usually destabilized or translationally inhibited in the cytoplasm. However, some Argonautes have nuclear localization and regulate gene expression on the transcriptional level. For instance, in *C. elegans*, NRDE-3 and HRDE-1 are nuclear Argonautes that silence genes on the transcriptional level in the soma and in the germline, respectively (Buckley et al., 2012; Guang et al., 2008).

*C. elegans*, like plants and yeast, has RNA-dependent RNA Polymerases (RdRPs) dedicated to the production of sRNAs. *C. elegans* has four RdRP genes, RRF-1/-2/-3 and EGO-1. It is believed that these RdRPs synthesize sRNA fragments in an unprimed manner (Billi et al., 2014). Two of these RdRPs, RRF-1 and EGO-1, generate sRNAs after target recognition by a primary Argonaute. These secondary sRNAs (22G-RNAs) contain a 5’-triphosphate group, have a bias for a 5’ guanosine and are mostly 22 nucleotides long (Billi et al., 2014). The RdRP enzyme RRF-3 is required for the biogenesis of another endogenous sRNA population, known as 26G-RNAs, which are mainly 26 nucleotides long, have a 5’ guanosine bias and a 5’-monophosphate (Conine et al., 2010; Gent et al., 2009, 2010; Han et al., 2009; Pavelec et al., 2009; Vasale et al., 2010). The fourth RdRP gene, RRF-2, has no described function in RNAi-related pathways.

26G-RNAs can associate with three Argonautes. During spermatogenesis, 26G-RNAs associate with the Argonautes ALG-3 and ALG-4 (from here on indicated as ALG-3/4). These Argonautes are required for normal fertility and mostly target spermatogenic transcripts, mediating posttranscriptional gene silencing (Conine et al., 2010, 2013; Han et al., 2009). Also, ALG-3/4 targets show a significant overlap with targets of CSR-1, an Argonaute protein that has been suggested to potentiate gene expression, rather than gene silencing (Conine et al., 2013). During oogenesis and embryogenesis, 26G-RNAs associate with the Argonaute ERGO-1 (Gent et al., 2010; Han et al., 2009; Vasale et al., 2010). In contrast to the ALG-3/4-bound 26G-RNAs, ERGO-1-bound 26G-RNAs are 2’Omethylated by HENN-1, which increases their stability (Billi et al., 2014). The main targets of ERGO-1 are recently duplicated paralogs and pseudogenes (Vasale et al., 2010). Upon target recognition, ERGO-1 triggers the production of 22G-RNAs. In turn, these 22G-RNAs direct gene-silencing and presumably associate with unknown cytoplasmic Argonautes, as well as the somatic nuclear Argonaute protein NRDE-3 (Guang et al., 2008; Vasale et al., 2010). NRDE-3 and other NRDE factors lead to transcriptional gene silencing of their targets, a process accompanied by H3K9 trimethylation of the target locus (Burkhart et al., 2011; Burton et al., 2011).

Mutants defective in the generation of 26G-RNAs, in particular those associated with ERGO-1, are hypersensitive to exogenous RNAi (exoRNAi). This enhanced RNAi (Eri) phenotype, is believed to stem from the fact that 26G-RNA pathways share common components with the exoRNAi pathway (Duchaine et al., 2006; Gu et al., 2009; Yigit et al., 2006). Interestingly, many of the identified proteins that restrict exoRNAi in wild-type animals form a complex: the ERI Complex (ERIC) (Duchaine et al., 2006; Thivierge et al., 2012; Yigit et al., 2006). ERIC has a core module that has been proposed to consist of the RdRP RRF-3 and its close interacting partners, the DExD/H box helicase DRH-3 and the Tudor domain-containing protein ERI-5 (Duchaine et al., 2006; Thivierge et al., 2012). To become active, this core complex needs to interact with DCR-1, an interaction that requires ERI-5 (Thivierge et al., 2012). Additionally, ERI-1 and ERI-3 are accessory factors of the ERIC that promote 26G-RNA biogenesis (Billi et al., 2014; Duchaine et al., 2006). Further mechanistic insights into ERIC assembly and function are severely lacking.

Besides 22G-and 26G-RNAs, *C. elegans* produces 21U-RNAs (Billi et al., 2014). The 21U-RNAs interact with PRG-1, one of the *C. elegans* Piwi protein homologs, and are also known as the piRNAs of *C. elegans* (Billi et al., 2014). In many organisms, the Piwi-piRNA pathway provides protection against TEs (Luteijn and Ketting, 2013), and also in *C. elegans*, 21U-RNAs contribute to the defense against TE activity (Bagijn et al., 2012; de Albuquerque et al., 2015; Phillips et al., 2015). Interestingly, 21U-RNAs can initiate a nuclear, 22G-RNA-mediated pathway. These 22G-RNAs, bound by the nuclear Argonaute HRDE-1 can affect histone modification patterns on targeted loci, and can establish a very stably inherited form of gene silencing (named RNA-induced epigenetic silencing or RNAe) that no longer depends on continued exposure to 21U-RNAs (Ashe et al., 2012; Buckley et al., 2012; Lee et al., 2012; Luteijn et al., 2012; Shirayama et al., 2012).

Genome-wide screens have uncovered many factors involved in the piRNA pathway and TE silencing in *Drosophila melanogaster* (Czech et al., 2013; Handler et al., 2013; Muerdter et al., 2013). Many of these factors are poorly conserved evolutionarily. Gametocyte-specific factor 1 (Gtsf1), a double CHHC zinc finger protein, represents one of the few Piwi pathway components that displays clear evolutionary conservation. dmGtsf1 is required for fertility and associates directly with Piwi (Dönertas et al., 2013; Ohtani et al., 2013). Interestingly, in the absence of Gtsf1, Piwi is still nuclear and loaded with piRNAs, but cannot silence TEs. Hence, dmGtsf-1 has been proposed to be required for the execution of Piwi-mediated silencing activities following target recognition. Also in mice, Gtsf1 is required for fertility and Gtsf1-related proteins have been shown to interact with Piwi proteins (Takemoto et al., 2016; Yoshimura et al., 2007, 2009, 2018).

The precise molecular function of GTSF1, or of its isolated domains is unknown. GTSF1 homologs have two tandem CHHC zinc finger domains and an unstructured C-terminal tail. *In silico* studies showed that CHHC zinc fingers are found in three protein families (Andreeva and Tidow, 2008): 1) U11-48K proteins, members of the alternative spliceosome, 2) TRM13 tRNA methyltransferases and 3) GTSF1-related proteins. These CHHC-domains behave as independent folding units and bind stoichiometrically to zinc (Andreeva and Tidow, 2008). The CHHC zinc finger of human U11-48K was shown to bind to the 5’ splice site of U12-dependent introns (Tidow et al., 2009), suggesting that CHHC zinc fingers bind RNA. Interestingly, the GTSF family is the only family of proteins that has two CHHC zinc fingers in tandem (Andreeva and Tidow, 2008).

Given its strong participation in Piwi-induced TE silencing in *Drosophila* and mouse, and that it is one of the few factors acting with piRNAs that displays wide conservation, we decided to characterize the function of GTSF-1 in *C. elegans*. Strikingly, we find that GTSF-1 is not involved in TE silencing and does not affect 21U-RNA production or activity in *C. elegans*. Instead, GTSF-1 associates with the RdRP RRF-3 and is required to assemble the ERI complex. We propose that GTSF1 proteins in general, may be present in smaller pre-complexes that may promote the assembly of larger protein-RNA complexes that elicit downstream enzymatic activities, such as sRNA production or the establishment of transcriptional silencing.

## Results

### GTSF-1 is enriched in the germline but not in P-granules

*T06A10.3*, the downstream partner of *lsy-13* in an operon on chromosome IV, was identified by reciprocal BLAST as the *C. elegans gtsf1* homolog, and was named *gtsf-1* (**Figure 1A**). GTSF-1, like its mouse and fly homologs, has two predicted CHHC zinc fingers (Andreeva and Tidow, 2008). The cysteine and histidine residues of the zinc fingers, as well as several acidic residues on the C-terminal region are conserved from worms and flies to mouse, zebrafish and human (**Figure S1A**). We produced three independent *gtsf-1* deletion alleles using CRISPR-Cas9 technology (Friedland et al., 2013) (**Figures 1A-B**, **S1B**). Five times outcrossed, homozygous *gtsf-1* mutants are fertile and do not show any obvious morphological defects. No GTSF-1 protein is detected in the mutants by Western blot, using an anti-GTSF-1 polyclonal antibody (**Figure 1C**). Expression of *lsy-13*, the operon partner, does not seem to be affected in *gtsf-1(xf43)* mutants (**Figure S1C**).

**Figure 1.**
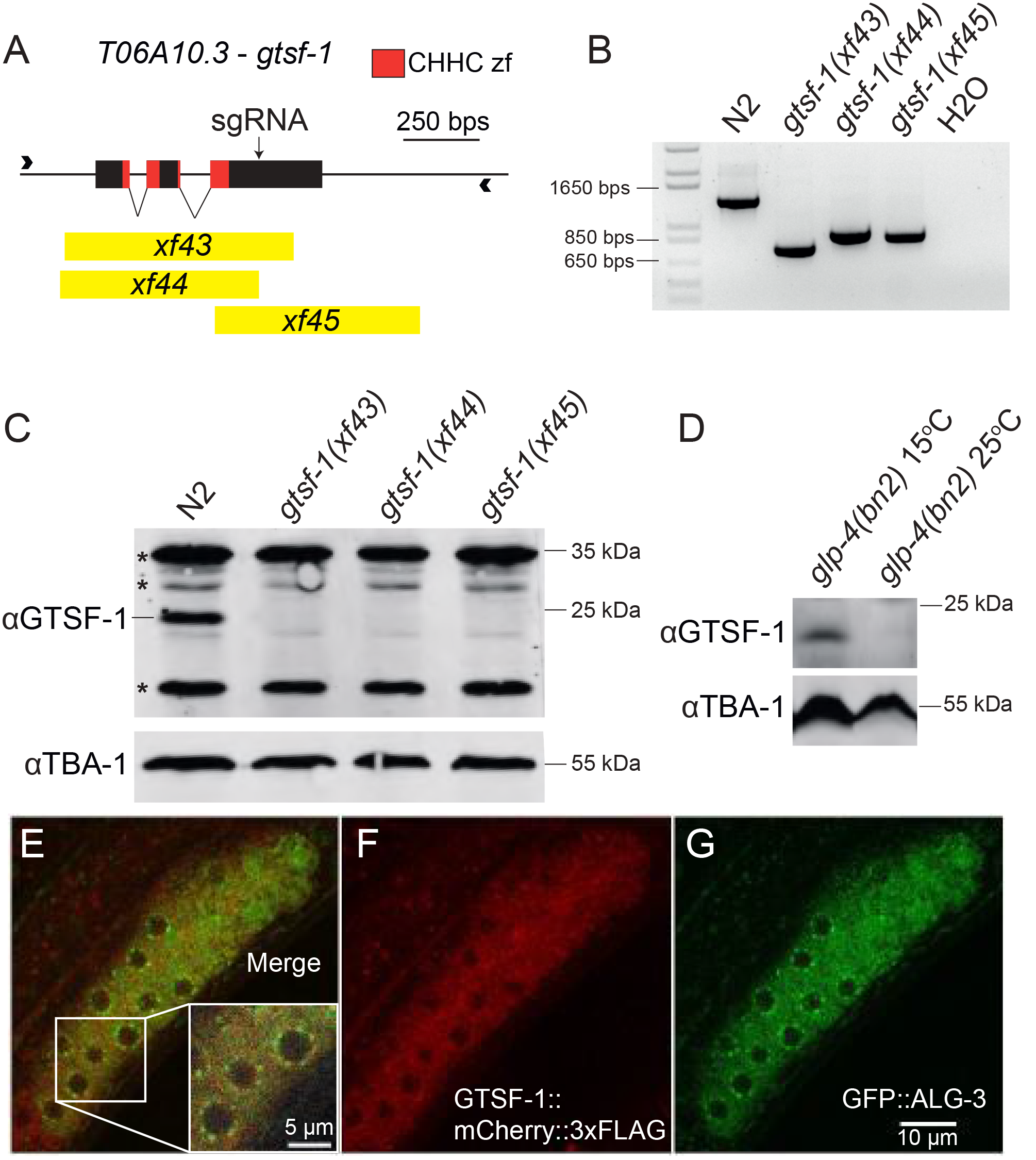
*T06A10.3*, the *C. elegans* homolog of *gtsf-1* is cytoplasmic and germline-enriched. (A) Overview of the *T06A10.3* gene in chromosome IV of *C. elegans*. The exons are represented as black boxes, the CHHC zinc finger domains are shown in red, and the black arrow corresponds to the cut site of the sgRNA used. The deletion alleles are represented in yellow. (B) PCR analysis of the deletion alleles using primers represented by arrowheads in (A). (C) Western blot analysis of mixed-stage wild-type and mutant worm extracts using a polyclonal anti-GTSF-1 antibody. TBA-1, one of the *C. elegans* alpha-tubulins was used as a loading control. (D) Western blot analysis of *glp-4(bn2)* mutant worms grown at the non-permissive temperature of 25°C, which precludes the development of the germline, and 15°C. (E-G) Representative confocal fluorescence microscopy images showing the presence of GTSF-1 and ALG-3 tagged proteins in a gonad of a L4 double transgenic worm, in the *alg-3/4; gtsf-1* triple mutant background. Scale bars correspond to 10 μm and 5 μm in the case of the inset.

To address the expression pattern of *gtsf-1* throughout development, we used publicly available RNA-sequencing datasets (Boeck et al., 2016). During embryonic development, larval development and adulthood, *gtsf-1* is moderately expressed (levels ranging from 0.4 to 7.2 depth of coverage per base per million reads [DCPM], **Figure S1D-E**). Notably, *gtsf-1* RNA levels are highest in the 4 cell stage and during the first 300 minutes of embryonic development (2.38-7.2 DCPM), suggesting that *gtsf-1* mRNA may be maternally deposited (**Figure S1D**). During larval development, *gtsf-1* mRNA reaches highest levels during the L4 and young adult stage (0.89-1.2 DCPM), correlating with germline development (**Figure S1E**).

To address potential germline enrichment of GTSF-1, we used *glp-4(bn2)* worms, which lack a germline when grown at 25°C. Western blot experiments on these animals (**Figure 1D**) indicate that GTSF-1 is enriched in the germline, since we could not detect GTSF-1 in *glp-4(bn2)* worms grown at 25°C. These data are supported by recent germline transcriptomes using dissected male and female gonads (Ortiz et al., 2014) that detected *gtsf-1* transcript in gonads irrespective of gender (**Figure S1F**). To address subcellular localization, we produced a *gtsf-1∷mCherry∷3xflag* single-copy transgene controlled by the germline-specific *gld-1* promoter (Merritt et al., 2008) and introduced it into a *gtsf-1(xf43)*; *alg-3(tm1155)*; *alg-4(ok1041)* triple mutant background, also expressing a GFP∷ALG-3 fusion protein. In these animals, we observed GTSF-1∷mCherry∷3xFLAG protein localized throughout the germline cytoplasm in L4 stage animals. GTSF-1 does not appear to be concentrated in P-granules, marked by GFP-tagged ALG-3 (**Figure 1E-G**), a known P-granule component (Conine et al., 2010).

These data indicate that *C. elegans* GTSF-1 is enriched in the germline cytoplasm, but mostly outside perinuclear granules.

### GTSF-1 is not involved in the 21U-RNA pathway and transposon silencing in *C. elegans*

Next, we wanted to address whether *gtsf-1* is involved in TE silencing. To test this, we used a strain with the *unc-22(st136)* allele, which has the *unc-22* gene interrupted by a Tc1 transposon (Ketting et al., 1999) (**Figure S2A**). Animals carrying the *unc-22(st136)* allele exhibit the so-called twitcher phenotype. When a gene that participates in TE silencing, such as *mut-7* (Ketting et al., 1999), is impaired in the *unc-22(st136)* background, TEs will become mobile and phenotypical reversions to wild-type movement can be observed. All three *gtsf-1* mutant alleles were crossed into the *unc-22(st136)* background and no reversions of the twitcher phenotype were observed after culturing the strains for several generations, in ten biological replicates per allele (comprising a reversion frequency of <10^−5^, **Figure S2B**).

To further characterize the role of *gtsf-1* in the sRNA pathways of *C. elegans*, we sequenced sRNAs from wild-type and *gtsf-1* synchronized gravid adults, in triplicates (experimental design in **Figure 2A**, sequencing statistics in **Supplemental Information**). To enrich for different sRNA species we employed different library preparations to each biological replicate. To increase the likelihood of cloning 22G-RNAs, which have a 5’ triphosphate, we used Tobacco Acid Phosphatase (TAP}. To enrich for sRNA species with a 2’-O methyl group on their 3’ end (21U-RNAs and ERGO-1-associated 26G-RNAs), we oxidized the RNA before library preparation with NaIO_4_ (the 2’-O-methyl group protects small RNAs from oxidation). Finally, we used untreated RNA to capture a higher fraction of sRNAs carrying a 5’ monophosphate, irrespective of their 3’ end methylation status (ERGO-1 and ALG-3/4-bound 26G-RNAs and miRNAs). The latter type of libraries will be hereafter referred to as “directly cloned”. “equences between 18-30 nucleotides were analyzed and read counts were normalized to the total number of mapped reads in each sample, excluding structural reads (see Methods).

**Figure 2.**
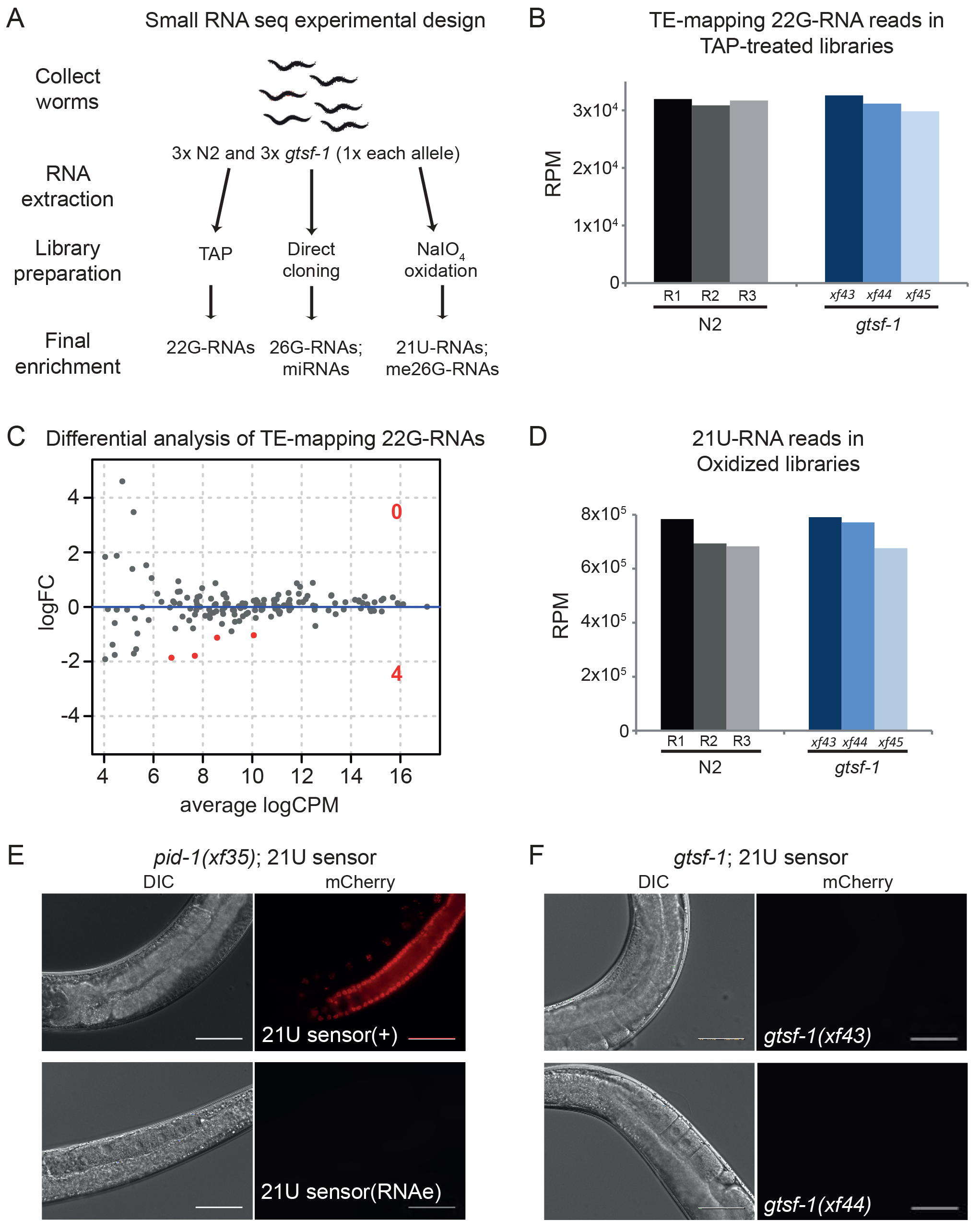
*C. elegans* GTSF-1 is not involved in the 21U-RNA pathway and TE silencing. (A) Experimental design of sRNA sequencing. Wild-type and *gtsf-1* mutant gravid adult worms were collected in triplicates. For *gtsf-1*, one sample of each allele was used as a biological replicate. Libraries were subjected to a triad of treatments to enrich for different small RNA species. TAP, Tobacco Acid Pyrophosphatase. (B) Similar abundance of TE-mapping 22G-RNA reads in TAP-treated libraries in wild-type (N2) and *gtsf-1* mutants (Welch two sample t-tests *p-value* = 0.75). Normalized levels in Reads Per Million (RPM) for each biological replicate are shown. (C) Differential analysis (MA-plot) of TE-mapping 22G-RNAs in *gtsf-1* mutants vs wild-type. sRNA reads from TAP-treated libraries were used for this analysis. Only four TEs show significantly downregulated (1% FDR) sRNA levels in *gtsf-1* mutants (see **Table S1**). LogFC, Log2 Fold Change. logCPM, log2 Counts Per Million. (D) Similar abundance of 21U-RNA reads in oxidized libraries in wild-type (N2) and *gtsf-1* mutants (Welch two sample t-tests *p-value* = 0.62). Normalized levels in Reads Per Million (RPM) for each biological replicate are shown. (E-F) Testing the participation of *gtsf-1* in the 21U-RNA pathway. For each figure, left panels are DIC while right panels show mCherry fluorescence channel. (E) Photomicrographs of adult worms carrying a 21U-RNA reporter in the *pid-1(xf35)* background. The panels above show a strain in which the 21U-sensor is still dependent on the 21U-RNA pathway, because in the absence of PID-1, mCherry can be observed in the germline. The panels below show a strain in which reporter silencing became independent of the 21U-RNA pathway, a state known as RNAe. (F) Micrographs of 21U-sensor;*gtsf-1* worms exhibiting the sensor repressed. This images are representative of 21U-sensor;*gtsf-1* worms originating from the crosses with both strains shown in (E) (schematics of the crosses are shown in Figure S2F,G). Scale bars represent 50 μm.

Consistent with the phenotypic experiments using the *unc-22(st136)* Tc1-transposition reporter, we did not observe major differences in sRNA reads mapping to TEs between wild-type and *gtsf-1* animals (**Figure 2B-C**, **Table S1**). Likewise, only two miRNAs were affected in *gtsf-1* mutants (**Figure S2C**, **Table S1**).

Also, the steady-state 21U-RNA levels are not significantly affected in *gtsf-1* mutants (**Figures 2D** **and S2D**). To further test participation of *gtsf-1* in the 21U-RNA pathway we performed crosses combining *gtsf-1* mutant alleles with an mCherry reporter for 21U-RNA activity (Bagijn et al., 2012; Luteijn et al., 2012) (**Figure S2E**). The reporter strains have a *pid-1(xf35)* mutation in the background to inform on the status of the sensor (**Figure 2E**) (Albuquerque et al., 2014), which can be under RNAe (insensitive to the presence of PID-1, **Figure 2E**, lower panels) or not (**Figure 2E**, upper panels). Loss of *gtsf-1* does not activate this reporter in either state, indicating it is not required for 21U-RNA-mediated silencing activity and RNAe (**Figures 2E-F** **and S2F-G**).

Overall, these data indicate that GTSF-1 is neither involved in TE silencing, nor in the 21U-RNA/RNAe pathway in *C. elegans,* in sharp contrast with the described function of Gtsf1 in mouse and fly.

### *gtsf-1* mutants recapitulate phenotypes of 26G-RNA pathway mutants

Given that *gtsf-1* is not involved in 21U-RNA-mediated gene silencing in *C. elegans*, we looked for other phenotypes that might be indicative of a role for GTSF-1 in other endogenous sRNA pathways. We noticed that populations of *gtsf-1* mutant animals grow slower compared to wild-type. This could reflect either developmental or fertility defects. When synchronized by bleaching, *gtsf-1* animals grew synchronous with wild-type (data not shown). In contrast, we noticed a striking reduction in brood size at 20°C, and temperature-sensitive sterility at 25°C (**Figure 3A**). When grown at 25°C, *gtsf-1* mutant animals mostly produced unfertilized oocytes (**Figures S3A-S3C**). At 20°C, both dead embryos and unfertilized oocytes were observed (data not shown). Importantly, two independent germline-specific *gtsf-1∷mCherry∷3xflag* transgenes (including *xfIs47*, the transgene shown in **Figures 1E-F**) completely rescue these defects (**Figures 3A** **and S3C**). These data clearly demonstrate that *gtsf-1* mutants display a temperature-sensitive fertility defect.

**Figure 3.**
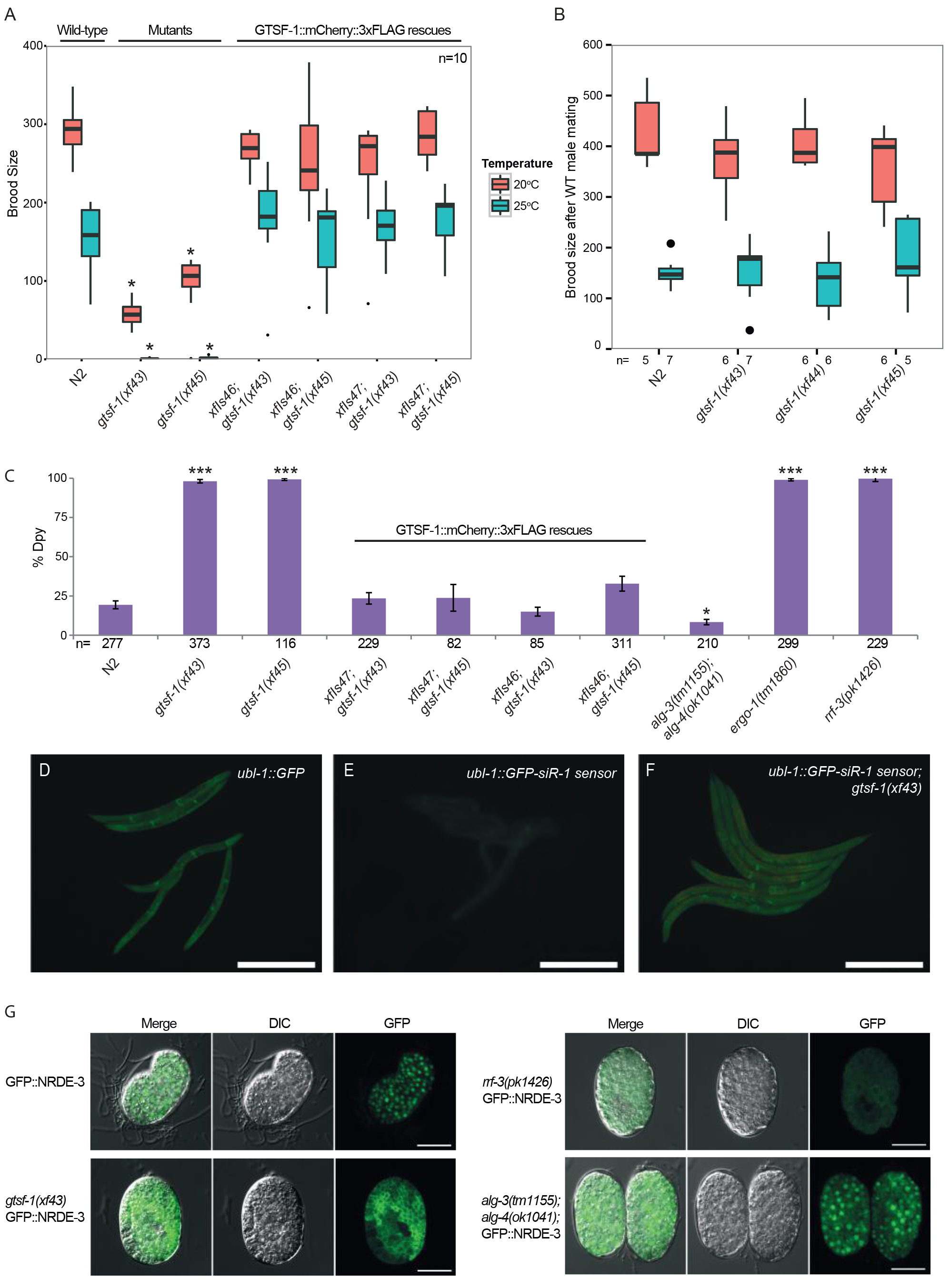
*gtsf-1* animals phenocopy 26G-RNA pathway mutants. (A) Boxplot of brood size counts at 20°C and 25°C. The progenies of 10 worms were counted for each strain and each temperature. Asterisks indicate *p-value*<0.0002 as assessed by Mann-Whitney-Wilcoxon tests comparing N2 worms with the other strains. Comparisons were done for each respective temperature. (B) Hermaphrodites with the genotypes indicated on the x-axis were mated with wild-type males and the progeny was counted. n for each condition is indicated in the figure below the x-axis. Mann-Whitney-Wilcoxon tests yielded *p-values*>0.4. (C) Assaying sensitivity to somatic *dpy-13* RNAi. The rescuing transgenes shown in **Figure 3A** are also assayed here. Total number of worms assayed is represented in the figure. Mann-Whitney-Wilcoxon tests were used to test if penetrance of *dpy-13* RNAi treatment was significantly different between N2 and mutant worms. Single asterisk indicates *p-value*=0.027 while triple asterisks indicate *p-values*<2.3e-05, *p-values* calculated using Mann-Whitney-Wilcoxon tests. Error bars represent the S.E.M. (D-F) GFP fluorescence images of worms carrying 22G-siR-1 sensor transgenes (see also Figure S3G). Scale bars represent 0.5 mm. (D) Animals carrying the control transgene with no 22G-siR-1 binding site. (E) Strains carrying the 22G-siR-1 sensor. (F) GFP signal in the absence of GTSF-1. (G) Micrographs of GFP∷NRDE-3 embryos in various genetic backgrounds. Scale bars represent 10 μm.

Temperature-sensitive sterility and embryonic lethality are recurring phenotypes of factors acting in endogenous sRNA pathways in *C. elegans*. For example, mutations in mutator genes, Eri genes, *rrf-3*, *drh-3* and *alg-3/4*, result in temperature-sensitive sterility at 25°C (Billi et al., 2014; Conine et al., 2010; Duchaine et al., 2006; Gent et al., 2009; Gu et al., 2009; Han et al., 2009; Ketting et al., 1999). In some of those mutants, like *alg-3/4, eri-1* and *rrf-3*, these fertility defects can be rescued by wild-type sperm, indicative of a sperm defect (Conine et al., 2010; Gent et al., 2009; Pavelec et al., 2009). Upon crossing *gtsf-1* hermaphrodites with wild-type males, both the reduced brood size at 20°C and the temperature-sensitive sterility at 25°C were rescued practically to wild-type levels (**Figure 3B**). Furthermore, we noticed that *gtsf-1* mutants have a mild high-incidence of males (him) phenotype (**Figure S3D**), again, similar to *alg-3/4*, many Eri and mutator mutants (Conine et al., 2010; Gent et al., 2009; Ketting et al., 1999).

One phenotype that distinguishes mutator mutants from Eri mutants is RNAi-sensitivity. Mutators are resistant to exogenous RNAi while Eri mutants are hypersensitive. *gtsf-1* mutants displayed normal sensitivity to RNAi against the germline gene *pos-1* (**Figure S3E**), but showed hypersensitivity to RNAi targeting somatic genes, as *dpy-13* (**Figures 3C** **and S3F**), *lir-1* and *pop-1* (data not shown), similarly to *rrf-3* and *ergo-1* mutant worms (Duchaine et al., 2006; Yigit et al., 2006). In contrast, *alg-3/4* double mutants did not display RNAi-hypersensitivity. Two independent, germline-specifically expressed *gtsf-1* transgenes rescued the RNAi hypersensitivity almost to wild-type levels (**Figures 3C** **and S3F**). We note that this rescue of a somatic phenotype with a germline-expressed transgene likely derives from the strong maternal effect of the 26G-RNA pathway (Zhuang and Hunter, 2011). We conclude that *gtsf-1* mutants have an Eri phenotype.

Loss of ERGO-1 and RRF-3, but not ALG-3/4, derepresses a ubiquitously expressed GFP transgene that reports on the activity of a specific 22G-RNA (**Figures 3D-E**, **S3G-H**), that is produced in response to ERGO-1 (Montgomery et al., 2012). GTSF-1 is also required for proper silencing of this transgene, indicating that the activity of GTSF-1 is required for ERGO-1/RRF-3-driven silencing (**Figures 3F** **and S3H**). We further tested GTSF-1 participation in the ERGO-1-dependent 26G-RNA pathway more broadly, by using a GFP∷NRDE-3 expressing transgene. GFP∷NRDE-3 in wild-type animals displays nuclear localization, but is cytoplasmic in *ergo-1* mutants because it fails to be loaded with 22G-RNAs (Guang et al., 2008). Nuclear localization is similarly affected by *gtsf-1(xf43)* and *rrf-3(pk1426)* mutation (**Figure 3G**). In contrast, *alg-3/4* mutations did not cause mislocalization of GFP∷NRDE-3 from the nucleus.

Overall, we conclude that *gtsf-1* mutants display phenotypes of *alg-3/4* and *ergo-1* mutants. As such, loss of GTSF-1 perfectly phenocopies loss of the RdRP enzyme RRF-3, suggesting that GTSF-1 acts at a very upstream step in the 26G-RNA pathway.

### 26G-RNA levels are strongly reduced in *gtsf-1* mutants

Given our phenotypic analysis, we reasoned that GTSF-1 may affect 26G-RNA biogenesis. Indeed, 26G-RNA levels are severely depleted in *gtsf-1* mutants (**Figure 4A**). This effect is observed both in the directly cloned as well as in the oxidized libraries, suggesting that both classes of 26G-RNAs, unmethylated (ALG-3/4-bound) and 2’O-methylated (ERGO-1-bound), respectively, are affected by GTSF-1 (**Figure 4A**). The levels of 26G-RNAs derived from all gene classes are similarly reduced upon loss of GTSF-1 (**Figure 4B**).

**Figure 4.**
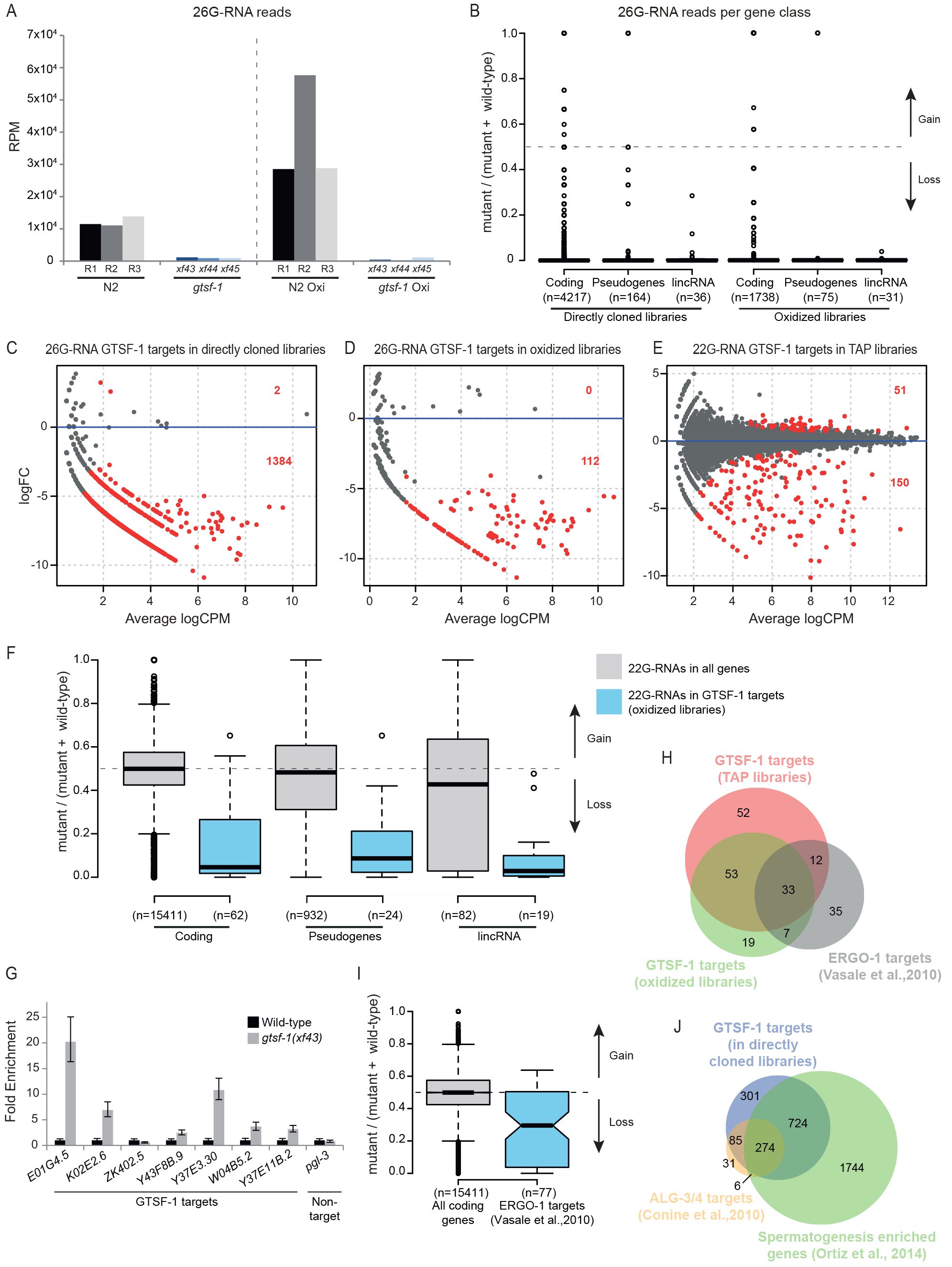
26G-RNAs are severely depleted in *gtsf-1* mutants. (A) Global levels of 26G-RNAs in wild-type and *gtsf-1* mutant worms, in RPM (Reads Per Million). Three biological replicates are shown, represented as R1-R3 for wild-type N2 worms. The dashed line separates the levels of 26G-RNAs in different library treatments: directly cloned libraries on the left, and oxidized libraries on the right. (B) Boxplot showing enrichment/depletion of normalized 26G-RNA reads per gene in *gtsf-1* mutants relative to wild-type, separated by gene class. All the genes in each class that had 26G-RNA mapped reads were used for this analysis. (C-E) Identification of GTSF-1 target genes that are significantly depleted of 26G-or 22G-RNA reads in the mutants in comparison to wild-type. Separate MA-plots are shown for the different library treatments. Statistically significant changes (1% FDR) are highlighted in red and their number is indicated. LogFC, Log2 Fold Change. LogCPM, log2 Counts Per Million. (F) Boxplot showing enrichment/depletion of 22G-RNA RNA reads in *gtsf-1* mutant in TAP libraries, by gene class, using all genes with mapped 22G-RNAs (grey boxes), and only 22G-RNAs that map to GTSF-1 targets (blue boxes), as defined in the oxidized libraries (D). (G) RT-qPCR of seven GTSF-1 targets and a non-target (*pgl-3*). Error bars represent the standard deviation of two biological replicates. *pmp-3* was used as the normalizing gene. (H) Venn diagram showing overlap of targets of the indicated libraries with previously defined ERGO-1 targets (Vasale et al., 2010). (I) Boxplot indicating enrichment/depletion of 22G-RNA levels (from the TAP-treated libraries) in all coding genes (grey box), and in ERGO-1 targets as defined by others. We used only 77/87 ERGO-1 RIP targets from Vasale et al, 2010, since for the remaining 10 targets, we did not have mapped reads. Notches represent the 95% confidence interval for each median. (J) Venn diagram showing overlap of targets of the indicated libraries with previously defined ALG-3/4 targets (Conine et al., 2010) and with genes enriched in the spermatogenic gonad (Ortiz et al., 2014).

Next, we defined high-confidence targets (at 1% FDR) of GTSF-1-dependent sRNAs for each library treatment (**Figure 4C-E,** lists of targets in **Table S1**). Targets were defined as genes that have a significant depletion of small RNAs in the mutant, in comparison with wild-type. The targets defined in the oxidized libraries (enriching for methylated 26G-RNAs) significantly overlapped with the targets of the TAP-treated libraries (enriching for 22G-RNAs, **Figure S4A**). These results suggest that genes that lose 2’-O-methylated 26G-RNAs also tend to lose downstream 22G-RNAs. This tendency is observed for all gene classes (**Figure 4F**). Next, we wanted to address if there are changes in GTSF-1 target gene expression concomitantly with loss of 26G-/22G-RNAs. Indeed, in the absence of GTSF-1, its targets are upregulated as assessed by RT-qPCR (**Figure 4G**, in levels consistent with previously published RT-qPCR data, see Duchaine et al., 2006; Pavelec et al., 2009; Vasale et al., 2010). Furthermore, our sets of GTSF-1 targets significantly overlap with a publicly available dataset from an ERGO-1 RIP (Vasale et al., 2010) (**Figure 4H**). Consistently, genes identified in the ERGO-1 RIP are depleted of 22G-RNAs in our TAP library dataset (**Figure 4I**). Of note, several of the GTSF-1 targets that were shown to be upregulated in **Figure 4G** were also identified as ERGO-1 targets (Vasale et al., 2010, namely *E01G4.5*, *K02E2.6*, *W04B5.2* and *Y37E11B.2*). ERGO-1 targets include paralog genes and pseudogenes (Vasale et al., 2010). Accordingly, we did not find strongly enriched gene ontology terms for the targets defined in the oxidized and TAP-treated libraries (**Table S1**). Furthermore, consistent with a role for GTSF-1 upstream of NRDE-3, we observed a significant overlap between GTSF-1-dependent sRNA targets and NRDE-3 targets (Zhou et al., 2014) (**Figure S4B**).

The 1384 targets defined by the directly cloned libraries (**Figure S4A**), extensively overlapped with ALG-3/4 targets as defined by small RNA sequencing of *alg-3/4* double mutants (Conine et al., 2010) (**Figure 4J**). Consistent with this, functional analysis for these 1384 GTSF-1 targets shows enrichment for sperm proteins, kinases and phosphatases (**Table S1**). As expected for ALG-3/4 targets, these GTSF-1-dependent loci extensively overlapped with spermatogenesis-specific genes as defined by others (Ortiz et al., 2014) (**Figure 4J**, **Table S1**).

To illustrate loss of small RNAs in *gtsf-1* mutants, exemplary genome tracks of GTSF-1 targets are shown in **Figure S4C-E**. Also, WormExp gene set enrichment analysis on GTSF-1 targets retrieved ERGO-1, ALG-3/4 and RRF-3 datasets, amongst many other datasets related to factors belonging to 26G-and 22G-RNA pathways (**Table S1**). Altogether, we conclude that both ALG-3/4-associated and ERGO-1-associated 26G-RNA populations, as well as the 22G-RNAs downstream of ERGO-1, are severely impacted by the loss of GTSF-1.

### GTSF-1 interacts with RRF-3

To identify interactors of GTSF-1 we performed Immunoprecipitation (IP) followed by label-free quantitative proteomics. IPs were performed in quadruplicates, in wild-type and *gtsf-1* mutant synchronized gravid adults using an anti-GTSF-1 antibody. Additionally, using an anti-FLAG antibody, we immunoprecipitated FLAG-tagged GTSF-1 from a strain carrying a rescuing transgene (the same used in **Figure 1E-F** and **Figure 3A,C**), using wild-type animals as a negative control. In both IP-mass spectrometry experiments, RRF-3 was the most enriched interactor (**Figure 5A-B**). Notably, in the transgene pull-downs (potentially an overexpression setup, because of the use of the *gld-1* promoter) we also observed slight enrichment of other known cofactors of RRF-3 in the 26G-RNA-producing ERIC (**Figure 5B**, represented by black dots). These IP experiments were also performed under more stringent wash conditions (600mM NaCl), in which case only the RRF-3 interaction was maintained (**Figure S5A-B**). We note that previous interactomics studies on Eri factors recovered GTSF-1 peptides, albeit with very low peptide coverage and without experiments addressing functionality (Duchaine et al., 2006; Thivierge et al., 2012). These observations support our results that GTSF-1 is associated with RRF-3 in the context of the ERIC.

**Figure 5.**
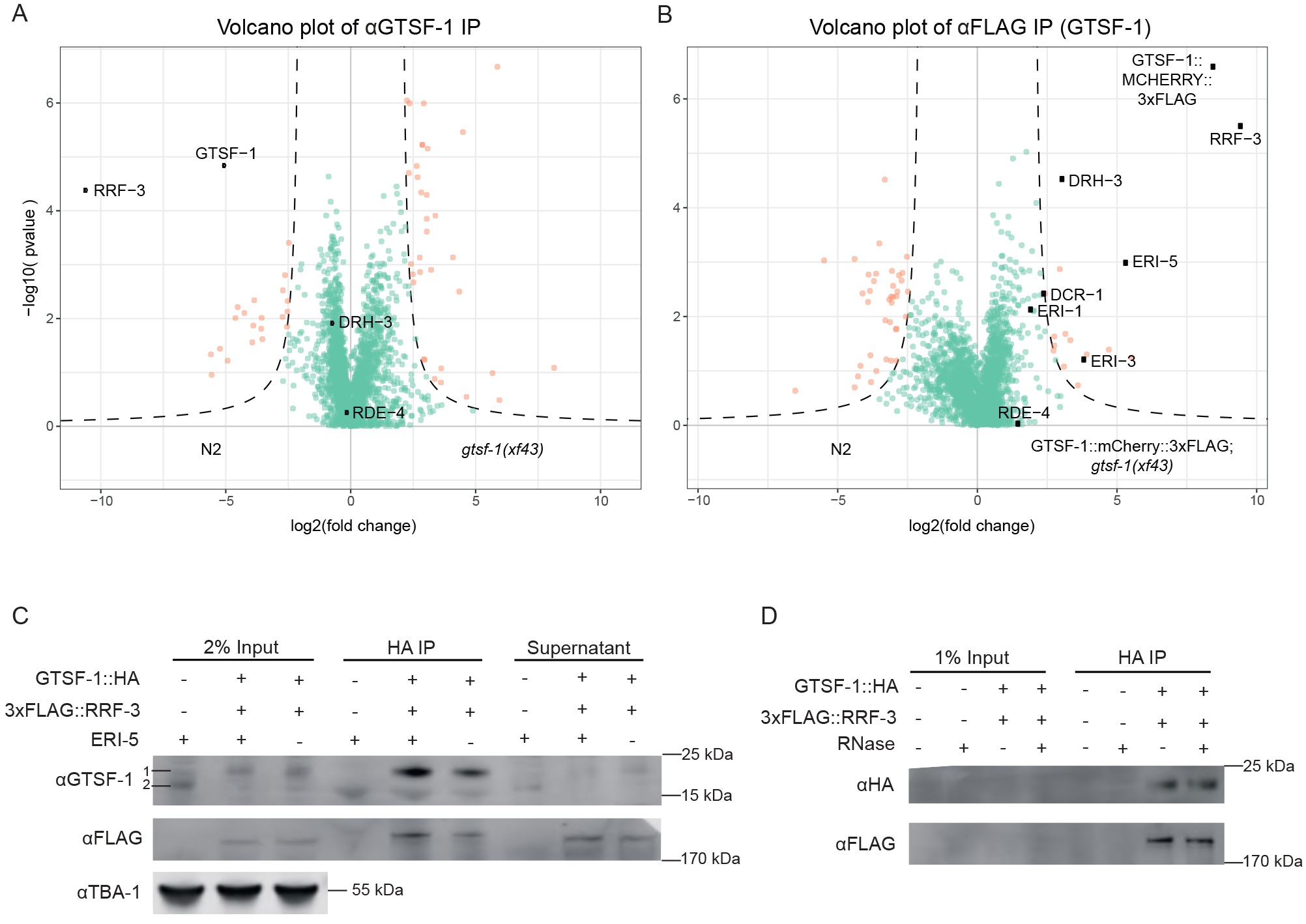
GTSF-1 interacts with RRF-3 in the adult germline, independently of RNA. (A-B) Volcano plots representing label-free proteomic quantification of GTSF-1 IPs from adult worm extracts. For each strain, IPs were performed and measured in quadruplicates. Log_2_ fold enrichment of individual proteins in one strain vs another is given on the x-axis. The y-axis indicates the Log_10_-transformed probability of the observed enrichments (see methods for details). Proteins in the background are represented as green dots while orange dots show enriched proteins. In (A) GTSF-1 was immunoprecipitated using our polyclonal anti-GTSF-1 antibody (in wild-type and *gtsf-1* mutant worms), while in (B) an anti-FLAG antibody was used to pull-down GTSF-1∷mCherry∷3xFLAG (in wild-type and strains carrying the rescuing transgene). (C) To test interaction between GTSF-1 and RRF-3 in adult worms by Western blot, GTSF-1∷HA was pulled-down via HA immunoprecipitation. Interaction was also tested in the presence/absence of ERI-5 by introducing an *eri-5(tm2528)* mutation in the background. Multi-channel secondary antibody detection was performed with an Odyssey CLx apparatus (see Methods). For the anti-GTSF-1, 1 represents GTSF-1∷HA and 2 represents untagged GTSF-1. (D) Testing RNA-dependency on the interaction between GTSF-1 and RRF-3 by adding RNase. Extracts from adult worms were used. Secondary antibody detection was performed with the Odyssey CLx setup.

To further characterize this interaction, we produced a single-copy transgene of 3xFLAG-tagged RRF-3. This transgene rescues the Eri phenotype and the fertility defects associated with loss of RRF-3 (**Figure S5C-D**), indicating it recapitulates wild-type RRF-3 function. We then used this transgene to validate the GSTF-1-RRF-3 interaction via Co-IP followed by Western Blot (**Figure 5C**). This interaction is not abrogated by RNase A treatment, indicating it is RNA-independent (**Figure 5D**).

These data clearly demonstrate that GTSF-1 interacts robustly with the RdRP enzyme RRF-3 and not with an Argonaute protein like its fly and mouse orthologs.

### The CHHC zinc fingers of GTSF-1 mediate the interaction with RRF-3

Next, we aimed to pinpoint the determinants of the GTSF-1/RRF-3 interaction. For this, we cloned and expressed GST-fused constructs with different GTSF-1 fragments (**Figure 6A-B**). Subsequently, we incubated these GST-fusions with embryonic extracts of a 3xFLAG∷RRF-3; *gtsf-1(xf43)*; *rrf-3(pk1426)* strain and pulled-down GST. Full length (FL) GTSF-1 pulls down 3xFLAG∷RRF-3 (**Figure 6B**), corroborating the results described above. The GST fusions to the individual zinc-fingers and the C-terminal tail did not pull-down 3xFLAG∷RRF-3 over background. Interestingly, when both CHHC zinc fingers are fused to GST, 3xFLAG∷RRF-3 can be efficiently retrieved (**Figure 6B**). None of the fusion proteins interacted with DCR-1 above background. We also created GST-GTSF-1 full length proteins with mutated zinc finger residues. Specifically, we mutated the cysteines of the zinc fingers to alanines (see **Figure S1A**). Notably, when we mutate the cysteines of individual zinc fingers, the interaction with 3xFLAG∷RRF-3 is slightly disturbed (**Figure 6C, see** Znf1-and Znf2-), but when all the four cysteines from both zinc fingers are simultaneously mutated, the interaction with 3xFLAG∷RRF-3 is abrogated (**Figure 6C,** see Znf12-). These results demonstrate that the zinc fingers of GTSF-1 are responsible for RRF-3 binding and suggest that both zinc fingers may act as a unit to mediate RRF-3 binding.

**Figure 6.**
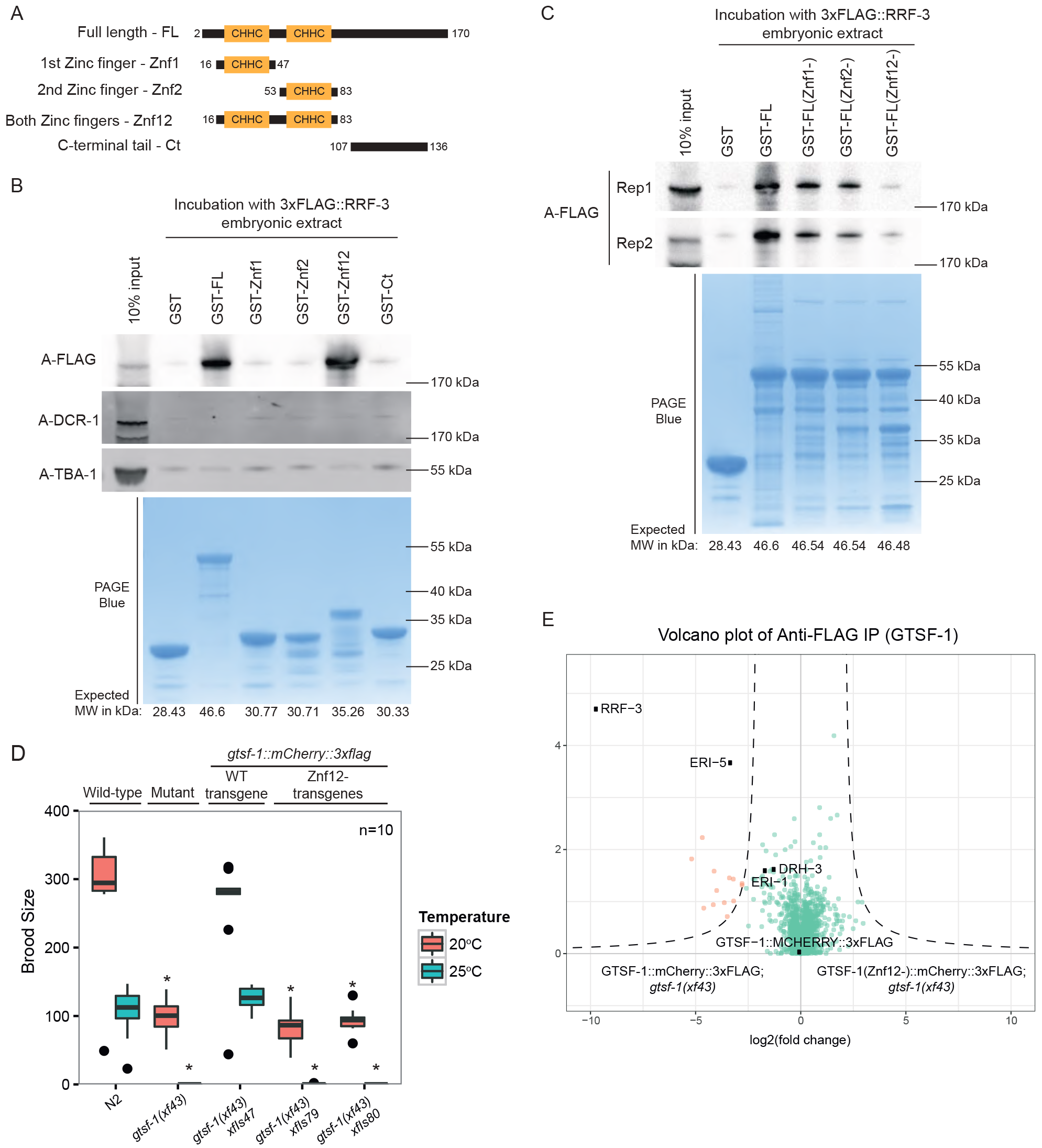
The tandem CHHC zinc fingers of GTSF-1 mediate the interaction with RRF-3. (A) Schematic representation of GST-fused GTSF-1 constructs produced for this study. Amino acid residues bordering the cloned regions are indicated in the figure. (B-C) Western blot analysis of GST-GTSF-1 pull-downs. (B) 5μg of GST-GTSF-1 fusion protein (conjugated with Sepharose GSH beads, see lower PAGE Blue panel) were each incubated with approximately 1 mg of total embryonic protein extract from a 3xFLAG∷RRF-3; *rrf-3(pk1426)*; *gtsf-1(xf43)* strain. 3xFLAG∷RRF-3 was detected using ECL while DCR-1 and TBA-1 were detected using the Odyssey CLx apparatus. (C) 5μg of various GST-GTSF-1 fusion proteins (conjugated with Sepharose GSH beads, see lower PAGE Blue panel), with the indicated cysteine to alanine mutations in the zinc fingers (1^st^ Zinc finger mutated – Znf1-; 2^nd^ zinc finger mutated – Znf2-; and both zinc fingers mutated-Znf12-) were each incubated with approximately 0.5 mg of total embryonic protein extract from a 3xFLAG∷RRF-3; *rrf-3(pk1426)*; *gtsf-1(xf43)* strain. 3xFLAG∷RRF-3 pull-down is shown for two independent biological replicates. (D) Brood size assay at 20°C and 25°C. The progenies of worms of the indicated genotype are plotted. The n for each sample is indicated in the x-axis. Asterisks indicate p-value<0.0037 as assessed by Mann-Whitney Wilcoxon tests comparing wild-type worms with the other strains. (E) Volcano plots showing label-free protein quantification of GTSF-1∷mCherry∷3xFLAG pull-downs. Pull-downs were performed in quadruplicate with adult worm extract. In (E), wild-type GTSF-1 fusion proteins are compared with GTSF-1 fusion proteins with zinc finger mutations (Znf12-). Proteins in the background are represented as green dots while orange dots show enriched proteins.

To address the *in vivo* relevance of the GTSF-1/RRF-3 interaction, we produced single-copy transgenes expressing GTSF-1 with mCherry and 3xFLAG tags, in which the CHHC cysteines in GTSF-1 were mutated to alanines (henceforth indicated as *gtsf-1*[*Znf12-*], see **Figure S1A**). Two independent *gtsf-1(znf12-)* transgene insertions do not rescue the Eri phenotype nor the fertility defects associated with GTSF-1, thereby phenocopying *gtsf-1* mutants (**Figure 6D** and **Figure S5D**). The lack of rescue is not due to poor expression of the (Znf12-) transgenes in the germline, although some degradation is observed (**Figure S5E**). Such partial degradation might be triggered by the disruption of the structural role that the zinc fingers have in GTSF-1. Moreover, subcellular localization of GTSF-1 is not affected by the zinc finger mutations (**Figure S5F**). FLAG pull-down followed by quantitative proteomics revealed that the GTSF-1(Znf12-) protein does not stably interact with RRF-3 (**Figure 6E**).

In the literature, several examples can be found of zinc fingers mediating both protein-protein and protein-nucleic acid interactions (Gamsjaeger et al., 2007). To address if GTSF-1 is interacting with RNA, we performed *in vitro* iCLIP (Sutandy et al., under review). We sought for a holistic approach, so we used *C. elegans* total RNA (rRNA-depleted) to test the binding of GTSF-1. Surprisingly, GTSF-1 was found not to crosslink with RNA above background levels (**Figure S5G**).

We conclude that GTSF-1 interacts with RRF-3 via its two tandem CHHC zinc fingers *in vitro* and *in vivo*. Since the GTSF-1/RRF-3 interaction is stable in presence of RNase (**Figure 5D**), and GTSF-1 does not seem to interact with RNA (**Figure S5G**), this suggests that the two CHHC zinc fingers in GTSF-1 act strictly as a protein-protein interaction domain.

### GTSF-1 is both in a precursor complex that is required for ERIC assembly, and in the mature ERIC

Previous studies on ERIC mostly focused on embryos (Duchaine et al., 2006; Thivierge et al., 2012). Next, we used embryonic extracts to probe the effect of GTSF-1 on ERIC. As in the adult germline, HA-tagged GTSF-1 pulls down 3xFLAG-tagged RRF-3 in embryos, as visualized by Western blot (**Figure 7A**). To circumvent potential overexpression (brought about by the transgene *gld-1* promoter), and to probe GTSF-1 interactions more broadly, we immunoprecipitated endogenous GTSF-1 and analyzed the precipitate with label-free quantitative mass spectrometry (**Figure 7B**). In this experiment, we observed a strong enrichment for RRF-3 and ERI-5, while all other known components of ERIC are either only mildly enriched, or not enriched at all, contrasting with the previously published molecular niche of RRF-3 in embryos (Duchaine et al., 2006; Thivierge et al., 2012).

**Figure 7.**
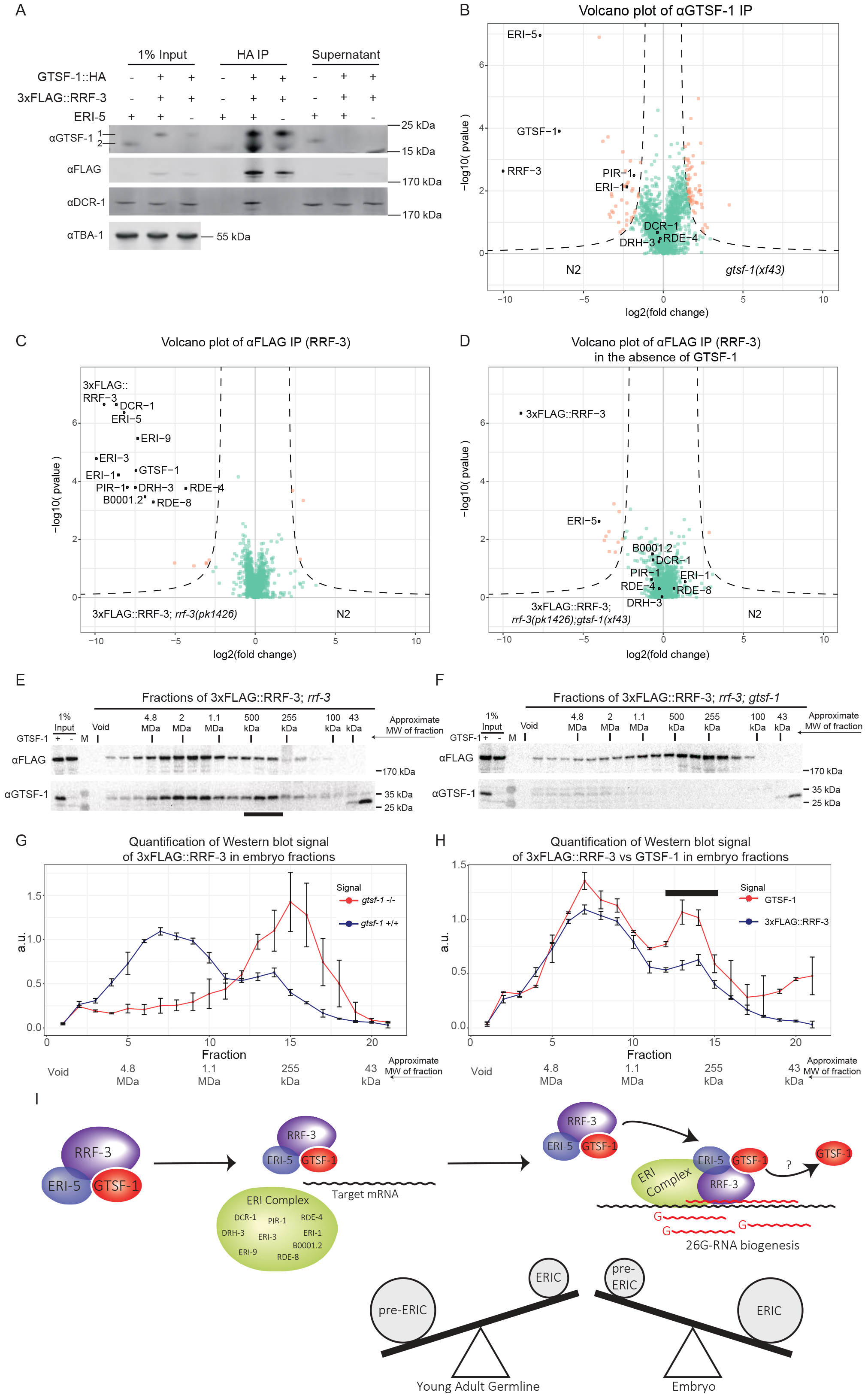
GTSF-1 is required for ERIC assembly. (A) Probing the interaction between GTSF-1 and RRF-3 by Western blot analysis, in embryonic extracts. GTSF-1∷HA was pulled-down via HA immunoprecipitation. Interaction was also tested in the presence/absence of ERI-5 by introducing an *eri-5(tm2528)* mutation in the background. Multi-channel secondary antibody detection was performed with an Odyssey CLx apparatus (see methods). (B) Label-free quantification of GTSF-1 IPs in embryos (comparing wild-type and *gtsf-1* mutant worms). IPs were done in quadruplicates, and a polyclonal anti-GTSF-1 antibody was used. (C-D) Volcano plots depicting quantitative proteomic analysis of RRF-3 pull-downs in the presence (C) and absence (D) of GTSF-1, in embryos. IPs were performed in quadruplicates. Proteins in the background are represented as green dots while orange dots show enriched proteins. (E-F) Size exclusion chromatography of 3xFLAG∷RRF-3-containing embryo extracts. Fractions were collected and probed for GTSF-1 and 3xFLAG∷RRF-3. Approximate Molecular Weight (MW) of the fractions is indicated. The calculation of these values according to protein standards is shown in the supplementary information. Fractions collected from extracts with GTSF-1 are shown in (E), and without GTSF-1 are shown in (F). (G-H) Comparison of size exclusion chromatography profiles of 3xFLAG∷RRF-3 and GTSF-1. Relative quantification was performed with the Western blot signal using ImageJ. Error bars represent standard deviation of two biological replicates. A.u., arbitrary units. (G) Comparison of profiles of 3xFLAG∷RRF-3 in the presence of GTSF-1 (blue line) and in the absence of GTSF-1 (red line). (H) Comparison of profiles of 3xFLAG∷RRF-3 (blue line) and GTSF-1 (red line). (I) A model for the function of GTSF-1. GTSF-1 forms a pre-ERI complex together with RRF-3 and ERI-5. GTSF-1 and ERI-5 are both required to incorporate RRF-3 into ERIC. Upon deposition of RRF-3 in ERIC, GTSF-1 may dissociate. See Discussion for details.

In order to test whether we can detect ERIC in our experimental setup, we performed IP-mass spectrometry on 3xFLAG∷RRF-3 from embryo extracts. This experiment clearly identified all known ERIC components (**Figure 7C**, black dots). In addition to the known ERIC components, we also found RDE-8 to strongly co-IP with RRF-3 under these conditions. RDE-8 and ERI-9, another previously identified ERIC factor (Thivierge et al., 2012), are paralog endonucleases that have been implicated in the 26G-RNA pathway (Gent et al., 2010; Tsai et al., 2015).

Given that the 3xFLAG∷RRF-3 IP results in the identification of ERIC in its entirety (**Figure 7C**), while the GTSF-1 IP retrieves only RRF-3 and ERI-5, we hypothesized that GTSF-1 binds non-ERIC-bound RRF-3. Is this non-ERIC-bound pool of RRF-3 perhaps a precursor complex that is required for ERIC formation? To test this, we performed a 3xFLAG∷RRF-3 IP in a *gtsf-1* mutant background, and again detected RRF-3 interactors through label-free quantitative mass spectrometry. Strikingly, in absence of GTSF-1, ERIC components no longer co-IP with RRF-3 (**Figure 7D**), with the sole exception of ERI-5. We then tested whether ERI-5 is required for interaction between GTSF-1 and RRF-3 and found that GTSF-1∷HA can still pull-down 3xFLAG∷RRF-3 in *eri-5* mutants (**Figures 5C and 7A**). Interestingly, we noticed that in the absence of ERI-5, both GTSF-1 and RRF-3 are partially destabilized in embryonic extracts (**Figure S6A**), while 3xFLAG∷RRF-3 is not destabilized in the absence of GTSF-1 (**Figure S6B**). These results suggest that 1) GTSF-1 is required to form mature ERIC from a RRF-3-ERI-5 precursor complex, where ERI-5 stabilizes RRF-3; 2) that GTSF-1 does not require ERI-5 to bind to RRF-3; and 3) ERI-5 does not require GTSF-1 to bind RRF-3.

To further test the idea that GTSF-1 is required to incorporate RRF-3 into ERIC, we performed size-exclusion chromatography with 3xFLAG∷RRF-3-containing embryonic extracts, followed by Western blot for GTSF-1 and FLAG. In wild-type embryos 3xFLAG∷RRF-3 displays a bimodal elution pattern. The main pool elutes in a broad range between 1-4 MDa, while a smaller fraction elutes at roughly 300-400kDa (**Figure 7E, G**). In absence of GTSF-1, 3xFLAG∷RRF-3 displays a single peak at roughly 250 kDa (**Figure 7F-G)**, consistent with RRF-3 bound to ERI-5 (61.6 kDa and 18.6 kDa are the predicted molecular weights for ERI-5 isoform A and B, respectively). These data support the hypothesis that GTSF-1 is required to incorporate an RRF-3/ERI-5 pre-complex into ERIC, via an RRF-3/ERI-5/GTSF-1 intermediate.

GTSF-1 and RRF-3 show very similar elution patterns (**Figure 7E, H**). This indicates that GTSF-1 remains within ERIC, at least for a significant time after its assembly. Results that we obtained by size-exclusion chromatography on young adult extracts are consistent with the embryo data: in young adults we also find that 3xFLAG∷RRF-3 and GTSF-1 display bimodal elution profiles (**Figure S6C-F**), with GTSF-1 again being essential to form ERIC (**Figure S6D-E**). Strikingly, both 3xFLAG∷RRF-3 and GTSF-1 show a more pronounced pre-ERIC peak when compared to embryos (compare **Figure S6C, F** with **Figure 7E, H**), suggesting ERIC assembly may be less active in the germline. Finally, both in embryos as well as in adults the ratio of pre-ERIC:ERIC is consistently higher for GTSF-1 than for RRF-3 (**Figures 7H** and **S6F**). This may indicate that GTSF-1 can dissociate from mature ERIC to form novel pre-ERIC complexes.

Taken together, these data show that GTSF-1 alternates between two states: one associated with the mature ERIC and another associated with an RRF-3 and ERI-5-containing pre-ERIC-complex. Also, and most importantly, this pre-complex is required to form a functional ERIC, competent for driving 26G-RNA biogenesis.

## Discussion

Here, we show that GTSF-1 does not participate in transposon silencing via the piRNA pathway in *C. elegans*, unlike GTSF-1 orthologs in flies and mice. However, like its orthologs, GTSF-1 is required for normal fertility. Surprisingly, GTSF-1 promotes 26G-RNA biogenesis by incorporating the 26G-RNA generating enzyme RRF-3 into a larger complex known as ERIC. GTSF-1 thus provides an enticing example of a conserved protein that achieves its function in sRNA pathways via different cofactors in different species, i.e. Argonaute proteins versus RdRP enzymes. Nevertheless, we propose that the function ascribed to *C. elegans* GTSF-1, of enabling the assembly of larger protein complexes from smaller subunits, may be evolutionarily conserved.

### The double CHHC zinc finger as a protein-protein interaction module

Typically, zinc fingers are known to mediate interactions with nucleic acid. Nevertheless, several cases were described in which zinc fingers mediate protein-protein interactions (Gamsjaeger et al., 2007). In some of these cases, zinc fingers of one protein interact directly with the zinc fingers of another protein (*e.g.* like GATA-1 and FOG)(Gamsjaeger et al., 2007).

We found that GTSF-1 interacts with RRF-3 via its tandem CHHC zinc fingers *in vitro* and *in vivo* (**Figure 6**). Interestingly, the zinc fingers individually could not interact with RRF-3 (**Figure 6B**). This suggests the two zinc fingers may function as one structural unit. Mutation of the cysteines of individual zinc fingers reduced but did not completely eliminate the interaction with RRF-3 (**Figure 6C**, see GST-GTSF-1 znf1-and znf2-). This could point at a certain structural robustness that allows one mutated zinc finger to fold relatively well when adjacent to a wild-type zinc finger. Of note, GTSF-1(znf12-) transgenes could not rescue *gtsf-1* mutant defects (**Figure 6D** and **S5D**), clearly showing that interaction with RRF-3 via its zinc fingers is key for GTSF-1 function *in vivo*.

These results differ from Piwi-Gtsf1 interaction data from *Drosophila* and mouse, in that the C-terminal tail (also referred to as “central region“} of dmGtsf1 was shown to interact with Piwi, and MIWI2 and MILI, respectively (Dönertas et al., 2013; Yoshimura et al., 2018). Also, GTSF-1 zinc finger mutants were still found to interact with Piwi in cell culture (Ohtani et al., 2013). We note, however, that 1) the zinc fingers of DmGTSF1 were not tested directly for interaction with Piwi, 2) the four cysteines of both zinc fingers were not simultaneously mutated, unlike our setup (**Figure 6**), and 3) consistent with our observations, zinc finger mutations are required for DmGtsf1 function, as assessed by transposon derepression (Dönertas et al., 2013; Ohtani et al., 2013). For Gtsf1l and Gtsf2, Gtsf1 paralogs in mouse, interaction with Piwi proteins and piRNA pathway cofactors was shown to be complex (Takemoto et al., 2016). For Gtsf1l, the double CHHC zinc fingers were shown, by *in vitro* GST pull-downs, to mediate interaction with MIWI and TDRD1. Interaction with MILI seems to be mediated by the “central region” encompassing the conserved acidic residues (**Figure S1A**). Conversely, GTSF2 interacts with MILI and TDRD1 via its CHHC zinc fingers, while it interacts with MIWI via its “central region”. It should be noted, however, that the relevance of these interactions has not been demonstrated *in vivo*. Nevertheless, it is possible that GTSF-1 may possess multiple interaction surfaces, with which it may be able to bring different complexes into close contact.

It seems that the CHHC zinc fingers present in GTSF proteins are not interacting with RNA, as was assumed after RNA-interaction was determined for the single CHHC zinc finger of U11-48K proteins (Andreeva and Tidow, 2008; Tidow et al., 2009). Interestingly, GTSF proteins are the only CHHC-containing protein family that has CHHC zinc fingers in tandem. It may be that this particular feature brought about structural possibilities that facilitate specific protein-protein interactions. We hypothesize that the tandem CHHC zinc fingers of GTSF1 homologs may generally function as one structural unit, with different structural characteristics than the individual U11-48K type CHHC zinc finger.

### A parallel between GTSF-1 in animals and Stc1 in fission yeast

In *S. pombe*, Stc1 is a protein that is required for sRNA-mediated centromeric heterochromatin formation (Bayne et al., 2010). More concretely, Stc1 bridges the Ago1 RNA-induced transcriptional silencing complex to the Clr4 methyltransferase complex. Although not phylogenetically related to GTSF-1 homologs, Stc1 has astonishingly similar structural features. It has an N-terminal LIM domain (which consists of two tandem zinc fingers) and a very acidic, unstructured, C-terminal domain, much like GTSF-1 (**Figure S1A**). Structure-function studies indicated that the tandem zinc fingers of Stc1 mediate a direct interaction with Ago1 while its C-terminal tail interacts with Clr4 (He et al., 2013). These modular protein-protein interactions nicely illustrate the bridging functions of Stc1.

In a similar fashion, *C. elegans* GTSF-1 may bridge RRF-3 and the rest of the ERIC. This would imply that the C-terminal tail of GTSF-1 would interact with another ERIC factor. We performed mass spectrometry of GST pull-downs of fusion constructs containing the C-terminal tail of GTSF-1. However, these experiments did not enrich for any ERIC factor, nor for any other plausible candidates (data not shown). It may be that this interaction is too transient to be detected in our experiments. The *in vitro* interaction studies of Gtsf1 proteins in mouse, described above, would also lend support to such a bridging function of GTSF1 in animals, i.e. reciprocally bridging MILI and MIWI complexes undergoing the ping-pong cycle.

Also in flies, GTSF1 might function to couple Piwi to downstream effector proteins such as Panoramix (Sienski et al., 2015; Yu et al., 2015). Possibly, this would need to be tested in specific developmental stages, since Piwi activity in flies was proposed to be primarily active in embryos (Akkouche et al., 2017).

GTSF1 homologs and Stc1 are not the sole examples of tandem zinc finger proteins with roles in small RNA pathways. A family of LIM-domain containing proteins in mammals was implicated in miRNA-mediated gene silencing (James et al., 2010). These LIM-domain proteins, LIMD1, Ajuba and WTIP, were found to bridge Ago1/2 with other factors, like eIF4E, in the molecular surroundings of the 5’ Cap structure. This mode of action will ultimately lead to translation inhibition of Ago1/2 targets (James et al., 2010). A more recent study has determined that the LIM domains of LIMD1 are the interaction surface with TNRC6A (Bridge et al., 2017). Moreover, LIMD1 bridges AGO2 to TNRC6A/miRISC (Bridge et al., 2017).

Altogether, it seems likely that small proteins with these structural modules, tandem zinc fingers and unstructured C-terminal domains, have convergently evolved as versatile bridges between different protein complexes with roles in small RNA pathways.

### How is the ERIC recruited to target RNA?

It is still unknown how ERIC is brought to, or assembled on target mRNA. How are the targets defined in the first place, and which ERIC component binds the RNA? To answer these questions, efforts should be made to identify the RNA-binding protein(s) involved in the recruitment of the mRNA. This could provide nice insights into the interplay between pre-ERIC (GTSF-1/ERI-5/RRF-3), ERIC and target mRNA.

We cannot fully exclude that the zinc fingers of GTSF-1, either together as a unit or individually, interact to some extent with RNA. However, the interaction with RRF-3 is not dependent on RNA (**Figure 5D**), and *in vitro* crosslinking experiments failed to show significant GTSF-1 association with RNA above background (**Figure S5G**). Hence, we believe GTSF-1 is unlikely to be responsible for RNA interaction during ERIC assembly.

Our FLAG∷RRF-3 pull-downs in embryos faithfully retrieved all known ERIC factors identified previously in other proteomics studies (Duchaine et al., 2006; Thivierge et al., 2012). Interestingly, we also retrieved one new RRF-3-interacting factor, RDE-8. This factor is a paralog of ERI-9 (Gent et al., 2010; Pavelec et al., 2009; Tsai et al., 2015), which was previously shown to interact with other ERIC factors (Thivierge et al., 2012). RDE-8 and ERI-9 are NYN ribonucleases, and have been previously shown to be involved in RNAi processes, including 26G-RNA biogenesis (Gent et al., 2010; Pavelec et al., 2009; Tsai et al., 2015). Their roles in 26G-RNA biogenesis seem to be independent of nucleic acid cleavage since: 1) ERI-9 lacks the conserved catalytic residues required for nucleic acid cleavage, and 2) RDE-8 transgenes with mutated catalytic residues still accumulate 26G-RNAs. Thus, it was proposed that RDE-8 and ERI-9 may have a structural role within the ERIC (Tsai et al., 2015). Alternatively, an attractive hypothesis is that RDE-8 and/or ERI-9 may be responsible for target mRNA recognition, or would play a role in stabilizing ERIC on its target RNA.

### What is the exact molecular function of GTSF-1?

We propose a model in which GTSF-1 and ERI-5 independently associate with RRF-3 to form a pre-ERIC (**Figure 7I**). This pre-ERIC is required to build a functional ERIC that drives 26G-RNA biogenesis. This process seems to be developmentally regulated, i.e. in the young adult germline there seems to be proportionally more GTSF-1/RRF-3 complex than in embryos. This means that this pre-complex may be “packaged” in the young adult germline to promptly initiate 26G-RNA biogenesis during embryonic development. Also, within the pre-ERIC, GTSF-1 and ERI-5 seem to have diverging roles. Both ERI-5 and GTSF-1 are required for building ERIC, but while ERI-5 seems to be required for the stability of GTSF-1 and RRF-3, GTSF-1 does not seem to be required for the stability of RRF-3.

Then, how does GTSF-1 exactly achieve its role? We consider a number of possibilities that are not mutually exclusive. First, GTSF-1 may be influencing the subcellular localization of RRF-3. Second, GTSF-1 may be chaperoning RRF-3 in a way that prompts conformational changes allowing RRF-3 to interact with other proteins. Third, GTSF-1 may allow RRF-3 to interact with target mRNA, which in turn may trigger ERIC assembly. In order to address these issues and fully understand how RRF-3 works, we will need to develop biochemical assays for ERIC assembly and function with purified components.

## Materials and Methods

### *C. elegans* genetics and culture

*C. elegans* was cultured on OP50 bacteria according to standard laboratory conditions (Brenner, 1974). Unless otherwise noted, worms were grown at 20°C. The Bristol strain N2 was used as the standard wild-type strain. The strains used and created in this study are listed in the **Supplemental Information**.

### Creation of *gtsf-1* mutants using CRISPR-Cas9 technology

*gtsf-1* mutant alleles were produced as described (Friedland et al., 2013). We successfully targeted the following sequencing on the third exon of *gtsf-1*: (GGAGCCGCTGGAGCTGAACG). Two other targeted sequences, cloned into p46169 in an identical fashion, did not yield any mutants, either alone or in combination: (GATAACATGCCCTTACAATT and GACGTCGGAAATCGAGAAAT).

N2 worms were injected with 150 ng/μl of Cas9 construct p46168 (a gift from John Calarco, Friedland *et al*., 2013), 135 ng/μl of sgRNA construct pRK1134 and 15 ng/μl of co-injection marker pCFJ104 (*Pmyo-3:mCherry:unc-54 3’UTR*, expresses mCherry in body wall muscle). F1 worms positive for mCherry expression in body wall muscle were isolated, allowed to self and then lysed in single worm lysis buffer (5 mM KCl, 2.5 mM MgCl_2_, 10mM Tris HCl pH=8.3, 0.45% NP40, 0.45% Tween20 and 0.01% gelatin). Subsequently, genotyping was performed with *Taq* Polymerase according to manufacturer’s instructions (New England BioLabs, M0273X). After isolation, *gtsf-1* mutant worms were outcrossed five times.

### Small RNA library preparation, sequencing and bioinformatics analysis

Detailed procedure for RNA isolation, small RNA enrichment, library preparation, bioinformatics analysis and sequencing statistics can be found in the **Supplemental Information**.

### Antibodies

Custom, affinity-purified rabbit anti-GTSF-1 antibodies were ordered from SDIX. The following protein sequence, comprising the last 91 amino acid residues of GTSF-1 (positions 79-169), was used as an antigen: (KRQSADLRRQLSLEPLELNVAEHLAAQKLRKEYEKDEESLDGSDDSDEDEEEKNLSVTSEIEKSDVEEVEMMLETINRLAYLEMKNDNLIL). The antibody (animal number Q5963) was used in a 1:500 dilution on Western blots and 2 μg were used on Immunoprecipitations. 2 μg of Anti-FLAG antibody (M2 clone, Sigma-Aldrich, F3165) were used on immunoprecipitations and a 1:5000 dilution was used for Western blot. DCR-1 antibody was a kind gift from Thomas Duchaine and it was used in Western blots in dilutions ranging from 1-3000 to 1-5000. More information on this antibody can be found elsewhere (Duchaine et al., 2006; Thivierge et al., 2012). A commercially available, mouse anti-tubulin monoclonal antibody (clone B-5-1-2, Sigma-Aldrich, T6074) was used in Western blots in a 1:10000 dilution to detect *C. elegans* TBA-1 as a loading control. A commercially available, rabbit anti-actin polyclonal antibody (Sigma-Aldrich, A5060) was used in Western blots in a 1:1000 dilution. 30 μL of suspension of E)view™ Red Anti-HA (mouse monoclonal antibody, clone HA-7) Affinity Gel (Sigma, E6779) were used for HA IPs. A mouse monoclonal Anti-HA antibody (clone HA-7, Sigma, H3663) was used in Western Blots with dilutions ranging from 1-500 to 1-1000.

### Mass Spectrometry

Details on worm sample collection and immunoprecipitation can be found in detail in the **Supplemental Information**. Immunoprecipitates were resuspended in NuPAGE LDS Sample Buffer 1X (Life technologies, NP0007) and 0.1 M DTT and heated at 70°C for 10 minutes. The respective samples were separated on a 4%-12% gradient Bis-Tris gel (NuPAGE Bis-Tris gels, 1.0 mm, 10 well, NP0321; Life Technologies) in 1x MOPS (NuPAGE 20x MOPS SDS running buffer, NP0001; Life Technologies) at 180 V for 10 min, afterwards separately processed by in-gel digest (Kappei et al., 2013; Shevchenko et al., 2007) and desalted using a C18 StageTip (Rappsilber et al., 2007).

The digested peptides were separated on a 25-cm reverse-phase capillary (75 μM inner diameter) packed with Reprosil C18 material (Dr. Maisch). Elution of the peptides was done along a 2h gradient from 2%-40% Buffer B (see Stage tip purification) with the EASY-nLC 1000 system (Thermo Scientific). Measurement was done on a Q Exactive Plus mass spectrometer (Thermo Scientific) operated with a Top10 data-dependent MS/MS acquisition method per full scan (Bluhm et al., 2016). The measurements were processed with the Max Quant software, version 1.5.2.8 (Cox and Mann, 2008) against the Uniprot *C. elegans* database (version of May, 2016) for quantitation. The mass spectrometry proteomics data have been deposited to the ProteomeXchange Consortium via the PRIDE partner repository with the dataset identifier PXD007665.

## Accession Numbers

Sequencing data have been deposited to the NCBI Gene Expression Omnibus (GEO) and proteomics data are available at the ProteomeXchange Consortium via PRIDE. GEO: GSE103432 [temporary token: qzgbqmqcjhabjml], PRIDE: PXD007665 [temporary username; temporary password: 4W5vdsva].

## Supplemental Information

Supplemental information includes supplemental experimental procedures, six figures and one table.

## Author Contributions

Conceptualization, M.V.A. and R.F.K.; Investigation, M.V.A., S.D., S.R., A.H., C.R.; Formal Analysis, M.V.A., S.D., S.R., E.K., A.H.; Visualization, M.V.A., S.D., S.R., E.K., A.H.; Writing – Original Draft, M.V.A. and R.F.K.; Writing – Review & Editing, all authors contributed; Funding acquisition, H.D.U., J.K., F.B., R.F.K.

## Acknowledgements

We thank all the members of the Ketting lab and Hans-Peter Wollscheid for great help and discussion. We thank Yasmin el Sherif, Svenja Hellmann and Isabel Pötzch for technical assistance. We acknowledge Chung-Ting Han and Hanna Lukas of the IMB Genomics core facility, the IMB Microscopy core facility and the IMB Media Lab. We are grateful to Thomas Duchaine and Ahilya Sawh for kindly providing the DCR-1 antibody and the *eri-5(tm2528)* strain. The authors thank Colin Conine and Craig C. Mello for providing the GFP∷ALG-3 strain. Also, the *Caenorhabditis* Genetics Center (CGC), which is funded by NIH Office of Research Infrastructure Programs (P40 OD010440), is acknowledged for providing worm strains. This work was supported by a Deutsche Forschungsgemeinschaft grant KE 1888/1-1 (Project Funding Programme to R.F.K.) and a ERC-AdG 323179 (to H.D.U.).

